# Compromising tyrosine hydroxylase function establishes a delusion-like temporal profile of reinforcement by dopamine neurons in *Drosophila*

**DOI:** 10.1101/2024.06.27.600982

**Authors:** Fatima Amin, Christian König, Jiajun Zhang, Liubov S. Kalinichenko, Svea Königsmann, Vivian Brunsberg, Thomas D. Riemensperger, Christian P. Müller, Bertram Gerber

## Abstract

For a proper representation of the causal structure of the world, one must consider both evidence for and evidence against causality. To take punishment as an example, the causality of a stimulus is reasonable if the stimulus precedes punishment, whereas causality can be ruled out if the punishment occurred first. This is reflected in the associative principle of timing-dependent valence reversal: aversive memories are formed when a stimulus occurs before the punishment, whereas memories of appetitive valence are observed when a stimulus is presented upon its relieving termination. We map the temporal profile of punishment induced by optogenetic activation of the PPL1-01 neuron in the fly *Drosophila melanogaster*, and find that impairment of tyrosine hydroxylase function, either acutely by pharmacological methods or by cell-specific RNAi, i) enhances learning with a time gap between stimulus and PPL1-01 punishment (trace conditioning), ii) impairs learning when the stimulus immediately precedes PPL1-01 punishment (delay conditioning), and iii) prevents learning about a stimulus presented after PPL1-01 punishment has ceased (relief conditioning). This implies a delusion-like state in which causality is attributed to cues that do not merit it (better trace conditioning), whereas both credible evidence for and credible evidence against causality is not properly appreciated (worse delay and relief conditioning). Under conditions of low dopamine, we furthermore observe a compensatory role for serotonin that is pronounced in trace conditioning, weaker in delay conditioning, and absent in relief conditioning. We discuss a disturbed dopamine-serotonin balance as an endophenotype for the positive and cognitive symptoms in schizophrenia.

## Introduction

A proper representation of the causal structure of the world is important for adaptive behavior. Indeed, distortions in the assignment of causes to effects can have consequences ranging from the comical to the lethal. Following early insight into the importance of ‘constantly conjoined events’ (Hume 1739-40/ 1978), causal learning is often studied in paradigms that vary the temporal relationship between cues and motivationally salient events (Shanks et al. 1989; Dickinson 2001). This may concern evidence in favor of a causal relationship with a punishment, for example, or evidence against such causality. That is, a cue X that has preceded a punishment is evidence for punishment, whereas it is evidence against such causation if the punishment came first and cue X followed it. Accordingly, in associative learning experiments, aversive memory for cue X is the result when X occurs before punishment, whereas a characteristically weaker and opposing, appetitive memory is the result when cue X is presented only upon the termination of punishment, at the moment of ‘relief’ (Solomon and Corbit 1974). Such timing-dependent valence reversal reflects a cross-species principle of reinforcement processing with broad implications in biomedicine and computational science (Malaka 1999; Gerber et al. 2014; Silver et al. 2016; Gerber et al. 2019).

In the fruit fly *Drosophila melanogaster*, timing-dependent valence reversal is mostly studied for the association between odor cues and electric shock punishment. After odor→ shock training the flies show learned avoidance of the odor, whereas learned approach is observed after shock→ odor training (Tanimoto et al. 2004). These memories are called punishment and relief memory, respectively (Gerber et al. 2014; Gerber et al. 2019). Punishment learning in *Drosophila* involves the coincidence of olfactory processing and shock-evoked dopaminergic reinforcement in the mushroom body, the highest brain center of insects (Heisenberg 2003; Cognigni et al. 2018; Li et al. 2020; Boto et al. 2020; Menzel 2022; Davis 2023). Pairings of odor presentation with the activation of the mushroom body input neuron PPL1-01 can establish aversive associative memory for the odor in a process that involves dopamine signaling from PPL1-01 to the mushroom body neurons (Aso et al. 2010; König et al. 2018; Aso et al. 2019) (synonyms for PPL1-01 are PPL1-γ1pedc and MB-MP1) (Figure 1A). However, although relief memory is observed for odors presented upon the termination of PPL1-01 activation (Aso and Rubin 2016; König et al. 2018), it is controversial whether this is mediated by dopamine, too. On the one hand, relief memory remained intact when we co-expressed in PPL1-01 both the optogenetic effector for activating it and an RNAi construct to knock-down the transcript for the tyrosine hydroxylase enzyme (TH) required for dopamine biosynthesis (König et al. 2018, loc. cit. figure 5). On the other hand, relief memory through PPL1-01 termination was abolished in loss-of-function mutants for TH (Aso et al. 2019, loc. cit. figure 5, supplementary figure 3C) (also see Handler et al. 2019). It is therefore imperative to clarify the contribution of dopamine in timing-dependent valence reversal by PPL1-01. A distinguishing feature of the present study is that we map out the full ‘fingerprint’ of PPL1-01 reinforcement across multiple temporal intervals (Figure 1B). Yet our results not only reconcile what appeared to be contradictory findings in König et al. (2018) and Aso et al. (2019). They further reveal a dissociation between two forms of punishment learning, namely for procedures with versus procedures without a time gap between odor presentation and PPL1-01 activation (‘trace’ versus ‘delay’ conditioning, respectively), moderated by both dopamine and serotonin. This unexpectedly complex modulation of reinforcement processing is discussed with respect to its implications for delusional causality attribution in psychosis.

**Figure 1:**
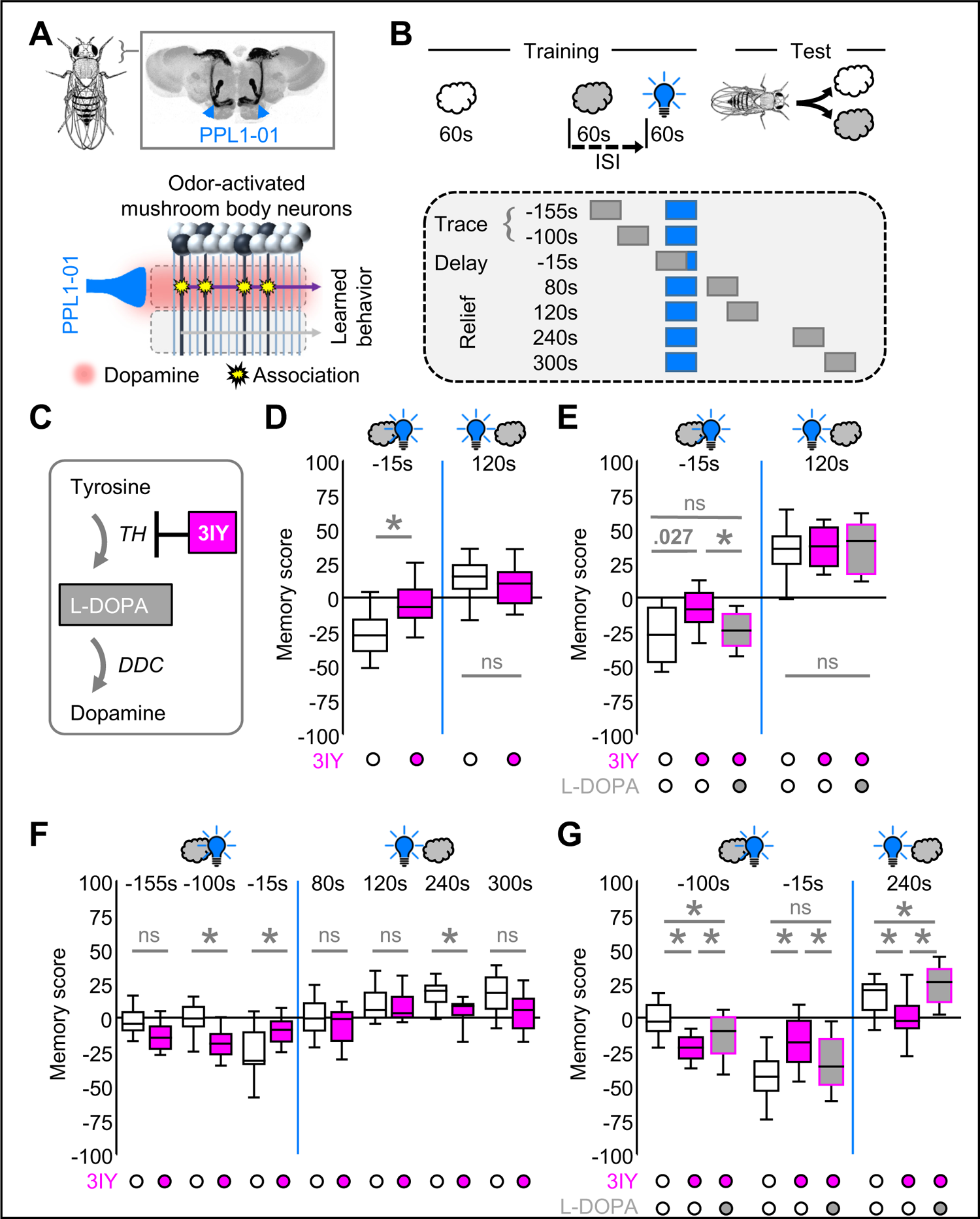
Pharmacologically inhibiting the TH enzyme both extends and blunts the temporal profile of reinforcement by PPL1-01. **(A)** Schematics of a fly, its brain, and the mushroom bodies (top) and a very highly simplified working hypothesis of association formation during punishment learning (bottom). The intrinsic neurons of the mushroom bodies represent odors in a sparse and combinatorial manner (mushroom body neurons in black). The dopaminergic PPL1-01 neuron (blue), which can be activated by e.g. electric shock punishment, intersects the axons of the mushroom body neurons in what is called the gamma1-peduncle compartment. Associative coincidence of odor activation and signaling from PPL1-01 (red shade within the gamma1-peduncle compartment) induces associative presynaptic plasticity (stars) at the synapses from the activated mushroom body neurons towards the output neuron of the gamma1-peduncle compartment (purple). This shifts the balance across the mushroom body output neurons to net avoidance as the learned behavior. In total the mushroom body has 15 compartments, one of which is sketched at the bottom. **(B)** Procedure for presenting the reference odor (open clouds), the paired odor (grey clouds), and optogenetic activation of PPL1-01 (blue light bulb). The interval between the onset of the paired odor and the onset of PPL1-01 activation is called the inter-stimulus-interval (ISI). For more details see Supplementary Figure 1. **(C)** Schematic of dopamine biosynthesis and of the inhibition of tyrosine hydroxylase (TH) by 3-iodo-L-tyrosine (3IY). The dopamine precursor 3,4-dihydroxy-L-phenylalanine (L-DOPA) should compensate for the effects of 3IY on dopamine levels. DDC: dopamine decarboxylase. Drug feeding was performed by the tissue paper method. **(D)** Relative to controls, punishment memory after odor→PPL1-01 training (ISI −15s) is decreased upon feeding of 3IY (N= 25, 25). Relief memory after PPL1-01→odor training (ISI 120s) is unaffected (N= 25, 23). **(E)** The decrease in punishment memory by 3IY can be rescued by additionally feeding L-DOPA (ISI - 15s) (N= 16, 16, 16). Relief memory is unaffected by 3IY, and by combining 3IY and L-DOPA (ISI 120s) (N= 16, 16, 15). **(F)** Mapping out the effect of 3IY on the temporal profile of PPL1-01 reinforcement (N= 20, 20; 20, 20; 19, 20; 20, 20; 20, 20; 20, 20; 20, 20). 3IY decreases punishment memory (ISI - 15s, delay conditioning) and leaves relief memory with an ISI of 120s unaffected. For a longer relief ISI of 240s a decrease in relief memory is revealed. For a training procedure with a −40s time gap between odor and PPL1-01 (ISI - 100s, trace conditioning), an increase in memory scores by 3IY is observed. **(G)** The effects of 3IY on memory scores after trace, delay, and relief conditioning (ISIs of −100s, −15s, and 240s, respectively) can be largely rescued, or even overcompensated, by L-DOPA (N= 30, 31, 31; 30, 31, 31; 30, 31, 30). Plotted in (D-G) are the memory scores according to equation 2, reflecting associative memory for the odor paired with optogenetic activation of PPL1-01; positive and negative memory scores reflect appetitive and aversive memory, respectively. Box plots represent the median as the middle line, 25%/75% quantiles as box boundaries, and 10%/90% quantiles as whiskers. Open box plots and circles refer to the control condition, magenta and grey fill to groups fed with 3IY or with 3IY plus L-DOPA, respectively. Flies were of the genotype PPL1-01>ChR-2XXL. * and “ns” indicate significance and non-significance, respectively, in MW-U tests at an error rate of 5%, adjusted according to Bonferroni-Holm, except for (E, ISI 120s) where “ns” indicates non-significance in a KW test. Data and statistical results are documented in Supplementary Table 1. The underlying preference scores are shown in Supplementary Figure 2. Memory scores separated by sex are shown in Supplementary Figure 3. The anatomical image of the mushroom body in (A) is modified from Heisenberg and Gerber (2008).

## Materials & Methods

### Fly strains

*Drosophila melanogaster* were reared in mass culture on standard food, at 60-70% relative humidity and 25°C, and under a 12h:12h light: dark cycle, unless mentioned otherwise. For the behavioral assays, 1-to-3-day-old adult flies were collected. Transgenic fly strains were used to express either the blue-light-gated cation channel ChR2-XXL, or both ChR2-XXL and an RNAi construct against the TH enzyme in the dopaminergic mushroom body input neuron PPL1-01. Specifically, males of the driver strain MB320C-split-GAL4 (covering the PPL1-01 neuron) (Bloomington stock center no. 68253; Aso et al. 2014) were crossed to females of the effector strains, which were either UAS-ChR2-XXL (Bloomington stock center no. 58374, Dawydow et al. 2014) or feature UAS-TH-RNAi in addition (Bloomington stock center no. 25796; Riemensperger et al. 2013). The flies from these crosses (henceforth PPL1-01>ChR2-XXL and PPL1-01>ChR2-XXL/TH-RNAi) were used for experiments and kept in darkness to avoid optogenetic activation by room light. Genetic controls carrying only the PPL1-01 driver or only the ChR2-XXL effector had previously been tested (König et al. 2018) and did not show memory upon pairing odor with blue light.

### Pharmacological manipulations

Unless mentioned otherwise, we used 3-iodo-L-tyrosine (3IY), an inhibitor of the TH enzyme which is rate-limiting for the synthesis of dopamine (Figure 1C), in a procedure that followed Thoener et al. (2021). Specifically, in different sets of newly hatched flies, either a plain 5% sucrose solution (CAS: 57-50-1, Hartenstein, Würzburg, Germany; in EVIAN water) was offered to the flies as their sole food, or it was offered in mixture with 5mg/ml 3IY (CAS: 70-78-0, Sigma, Steinheim, Germany; stored at −20°C) or in mixture with 5mg/ml 3IY plus 10mg/ml 3,4-dihydroxy-L-phenylalanine (L-DOPA), a precursor of dopamine (CAS: 59-92-7, Sigma, Steinheim, Germany). Mixtures were prepared by a shaker at high speed for approximately 60min. Specifically, flies were transferred to small plastic vials (diameter: 25mm; height: 60mm; volume: 30ml; K-TK, Retzstadt, Germany) with tissue paper (Fripa, Düren, Germany) soaked with 1.8ml of the solutions mentioned above, kept at 25°C and used for experiments after 36-40h. This procedure is henceforth called the tissue paper method.

We used para-chlorophenylalanine (PCPA, a.k.a. DL-4 Chlorophenylalanine, fenclonine) (CAS: 7424-00-2, ThermoFischer, Acros Organics, Geel, Belgium; stored at 4°C), an inhibitor of the enzyme tryptophan hydroxylase (TPH), which is rate-limiting for serotonin synthesis, in a procedure that followed Pooryasin and Fiala (2015). Specifically, newly hatched flies were transferred to large plastic vials (diameter: 46mm; height: 102mm; volume: 170ml; K-TK, Retzstadt, Germany) with wet tissue paper and starved for 48h at 18°C. After starvation, separate sets of flies were either transferred to small vials containing 1ml of freshly prepared standard food medium mixed with 200µl of 5% sucrose solution and 200µl of water (EVIAN) and left with this as their sole food, or they were kept with this mixture plus in addition either 1.25mg/ml of 3IY, or 1.25mg/ml of 3IY plus 60mg/ml of PCPA, or 60mg/ml of PCPA. In all cases, the flies were kept at 25°C room temperature and used for experiment 4 days later. Mixtures were prepared by a shaker at high speed for approximately 30min. This procedure is henceforth called the food method.

### Behavioral experiments

Behavioral experiments for the association of odor with the optogenetic activation of PPL1-01 followed König et al. (2018), unless mentioned otherwise. In brief, these experiments took place in a custom-made set-up (CON-ELEKTRONIK, Greussenheim, Germany) (modified from Tully and Quinn, 1985) that allowed the simultaneous handling of four groups of flies, each with 30-40 flies. During training, dim red light was used to allow minimal vision for the experimenter while the ChR2-XXL channels remained mostly closed. Blue light for opening the ChR2-XXL channels and thus for neuronal activation was turned on only briefly and in the temporal relationship to odor presentation as described below and throughout the Results section. In all cases, blue light was presented in a pulsatile manner as 12 pulses, each 1.2s long and followed by the next pulse with a 5s onset-to-onset interval. Once the training had concluded, the testing was carried out in darkness. As odorants, 50µl benzaldehyde (BA) and 250µl 3-octanol (OCT) (CAS 100-52-7, 589-98-0; Fluka, Steinheim, Germany) were applied to 1cm-deep Teflon containers of 5mm or 14mm diameter, respectively.

The key variable for the behavioral experiments is the relative timing (or inter-stimulus-interval, ISI) of the pairing between an odor and the optogenetic activation of PPL1-01 by blue light (Figure 1B, Supplementary Figure 1). The ISI is defined as the time difference between the onset of the blue light and the onset of the paired odor presentation. In principle, either the paired odor was presented first followed by blue light stimulation (forward conditioning, defined as negative ISIs), or the blue light was presented first followed by presentation of the paired odor (backward conditioning, positive ISIs). Specifically, separate groups of flies were trained with either of seven ISIs (−155s, −100s, −15s, 80s, 120s, 240s, 300s). Given the 60s-duration of blue light stimulation and the 60s-duration of the paired odor presentation, this resulted in gaps between the two events of −95s and −40s for the two longest forward conditioning ISIs (−155s, −100s) (‘trace’ conditioning), in a partial overlap for the −15s ISI (‘delay’ conditioning), and in gaps of 20s to 240s for the backward conditioning ISIs (‘relief’ conditioning). In all cases, a second odor was presented as a reference, explicitly unpaired from blue light during training. The use of BA and OCT as the paired and the reference odor was balanced across repetitions of the experiment.

After one cycle of training, which lasted for a total of 15min, the flies were given a 4min accommodation period and were then shifted to the choice point of a T-maze apparatus, with the paired and the reference odor on either side. After 2min, the arms of the T-maze were closed and the numbers of flies (#, as the sum of male and female flies) in each arm was counted by an assistant blind to the experimental conditions to calculate the preference for BA as:

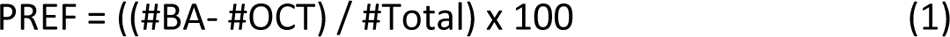

Positive PREF scores thus indicate preference for BA over OCT, and negative scores indicate preference for OCT over BA. From these scores, taken after either BA or OCT had been paired with PPL1-01 activation (BA+, or OCT+, respectively), an associative memory score was calculated to average out odor-specific and non-associative effects as:

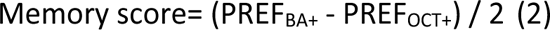

Negative memory scores thus indicate conditioned avoidance of the paired odor, and positive scores indicate conditioned approach to it.

To quantify the effect of compromising TH function on memory scores, the difference between the memory scores of the TH-compromised condition and the memory scores of the control condition was determined.

### Measurement of biogenic amine levels

After drug feeding using the tissue paper method, we performed a high-performance liquid chromatography (HPLC) analysis of brain-wide biogenic amine levels. For each of the respective treatments, 6 male and 6 female brains were dissected in Ca^2+^-free saline and, separated by sex, immediately frozen in −80°C liquid nitrogen. The samples were analyzed using HPLC with electrochemical detection to measure dopamine and serotonin levels. The column was an ET 125/2, Nucleosil 120-5, C-18 reversed phase column (Macherey & Nagel, Düren, Germany). The mobile phase consisted of 75mM NaH2PO4, 4mM KCl, 20μM EDTA, 1.5mM sodium dodecyl sulfate, 100μl/l diethylamine, 12% alcohol and 12% acetonitrile, adjusted to pH 6.0 using phosphoric acid (Carl Roth GmbH, Karlsruhe, Germany). The electrochemical detector (Intro) (Antec, Alphen, The Netherlands) was set at 500 mV versus an ISAAC reference electrode (Antec, Alphen, The Netherlands) at 30°C. This setup allows the simultaneous measurement of dopamine and serotonin (Amato et al. 2020; Kalinichenko et al. 2021).

### Statistical analyses

We used non-parametric statistical tests throughout (Statistica 11.0; StatSoft Hamburg, Germany, and R 2.15.1, www.r-project.org). For comparisons across more than two groups, the Kruskal-Wallis test (KW) was applied. For subsequent pair-wise comparisons between groups, Mann-Whitney U-tests (MW-U) were performed. To test whether values of a given group differed from chance levels, i.e., from zero, one-sample sign tests (OSS) were used. When multiple tests of the same kind were performed within one experiment, significance levels were adjusted by a Bonferroni-Holm correction to keep the experiment-wide type 1 error limited to 0.05. Data are presented as box plots which represent the median as the middle line and the 25%/75% and 10%/90% quantiles as box boundaries and whiskers, respectively. Data and statistical results are documented in the Supplementary Table 1.

## Results

### Pharmacological inhibition of TH both extends and blunts the temporal profile of PPL1-01 reinforcement

Flies expressing the blue-light-gated ion channel ChR2-XXL for optogenetic activation of the PPL1-01 neuron showed punishment memory after odor→PPL1-01 training (Figure 1D; ISI - 15s). Acute feeding of 3IY, an inhibitor of the TH enzyme required for dopamine synthesis, impaired such punishment memory (Figure 1D; ISI - 15s) (for a repetition see Supplementary Figure 4). Of note is that 3IY feeding leaves task-relevant sensory-motor faculties intact (Thoener et al. 2021). In a further repetition of the experiment the effect of 3IY feeding on punishment memory could be rescued by an additional feeding of L-DOPA (Figure 1E; ISI - 15s). In contrast, relief memory after PPL1-01→odor training was unaffected by feeding of 3IY (Figure 1D, Figure 1E; ISI 120s).

We next mapped out the effect of 3IY feeding on the temporal profile of PPL1-01 reinforcement more systematically, that is on the association of odor and PPL1-01 activation across multiple intervals between these events. For both the intervals used before, this once more replicated the finding that 3IY feeding leads to a decrease in punishment memory, and that there is a lack of such a detrimental effect on relief memory (Figure 1F; ISIs of −15s and 120s, respectively). Strikingly, however, 3IY feeding increased (sic) punishment memory when there was a −40s gap between the offset of the odor and the start of PPL1-01 activation (Figure 1F; ISI - 100s), and decreased relief memory for relatively long intervals between PPL1-01 activation and odor presentation (Figure 1F; ISI 240s, corresponding to a 180s gap). In a follow-up experiment we confirmed these three kinds of effect exerted by 3IY feeding and showed that they can be rescued by additionally feeding L-DOPA (Figure 1G). HPLC measurements of whole-brain homogenates upon 3IY feeding reveal a selective decrease in dopamine but not in serotonin levels, which was likewise rescued by additionally feeding L-DOPA (Figure 2).

**Figure 2:**
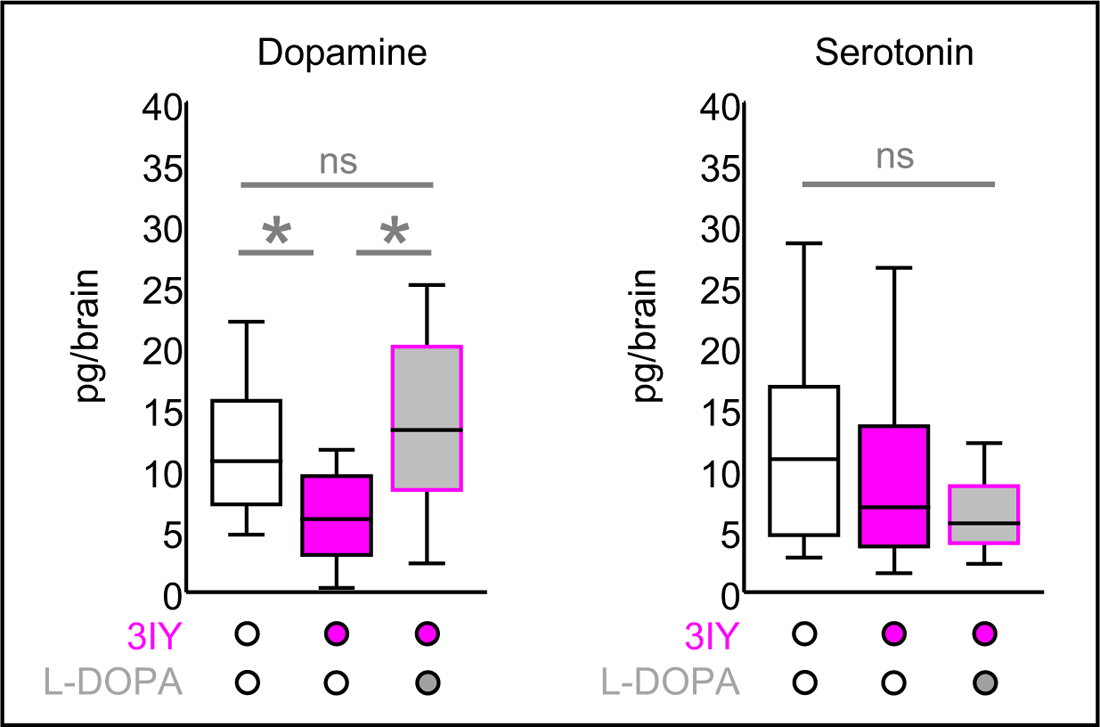
Pharmacologically inhibiting the TH enzyme reduces brain levels of dopamine Whole-brain levels of dopamine and serotonin after feeding 3-iodo-L-tyrosine (3IY), an inhibitor of the tyrosine hydroxylase (TH) required for dopamine biosynthesis. Feeding of 3IY reduced dopamine levels, an effect that was restored by feeding the dopamine precursor 3,4-dihydroxy-L-phenylalanine (L-DOPA) in addition (N= 20, 20, 20). Drug feeding was without effect on serotonin levels (N= 20, 20, 20). It was performed by the tissue paper method. Box plots represent the median as the middle line, 25%/75% quantiles as box boundaries, and 10%/90% quantiles as whiskers. Open box plots and circles refer to the control condition, magenta and grey fill to groups fed with 3IY or with 3IY plus L-DOPA, respectively. Flies were of the genotype PPL1-01>ChR-2XXL. * indicates significance in MW-U tests at an error rate of 5%, adjusted according to Bonferroni-Holm. “ns” indicates non-significance in such a MW-U test (dopamine) or in a KW test (serotonin). Data and statistical results are documented in Supplementary Table 1. Data separated by sex are shown in Supplementary Figure 5.

These results suggest that optogenetic activation of PPL1-01 establishes both punishment memory and relief memory through a 3IY-sensitive, TH-dependent, dopaminergic process. To our surprise, we notably found that the compromising of TH function had opposite effects upon training with a −40s gap between odor and PPL1-01 activation (ISI - 100s, punishment memory after trace conditioning) from training without such a gap (ISI - 15s, punishment memory after delay conditioning) (Figure 1F, Figure 1G).

### Local knock-down of TH likewise both extends and blunts the temporal profile of PPL1-01 reinforcement

We next asked whether the three observed effects of compromising TH function on memory scores, namely on punishment memory after i) trace and ii) delay conditioning, as well as on iii) relief memory, relate to dopaminergic signaling from PPL1-01 itself, rather than from dopaminergic neurons downstream of PPL1-01. We therefore co-expressed in PPL1-01 both the optogenetic effector and an RNAi construct for the knock-down of the TH enzyme (Riemensperger et al. 2013) and mapped out the temporal profile of PPL1-01 reinforcement. This again revealed an increase (sic) in punishment memory after trace conditioning (ISI - 100s), a decrease in punishment memory after delay conditioning (ISI - 15s), as well as a decrease in relief memory (ISI 300s) (Figure 3A, Figure 3B).

**Figure 3:**
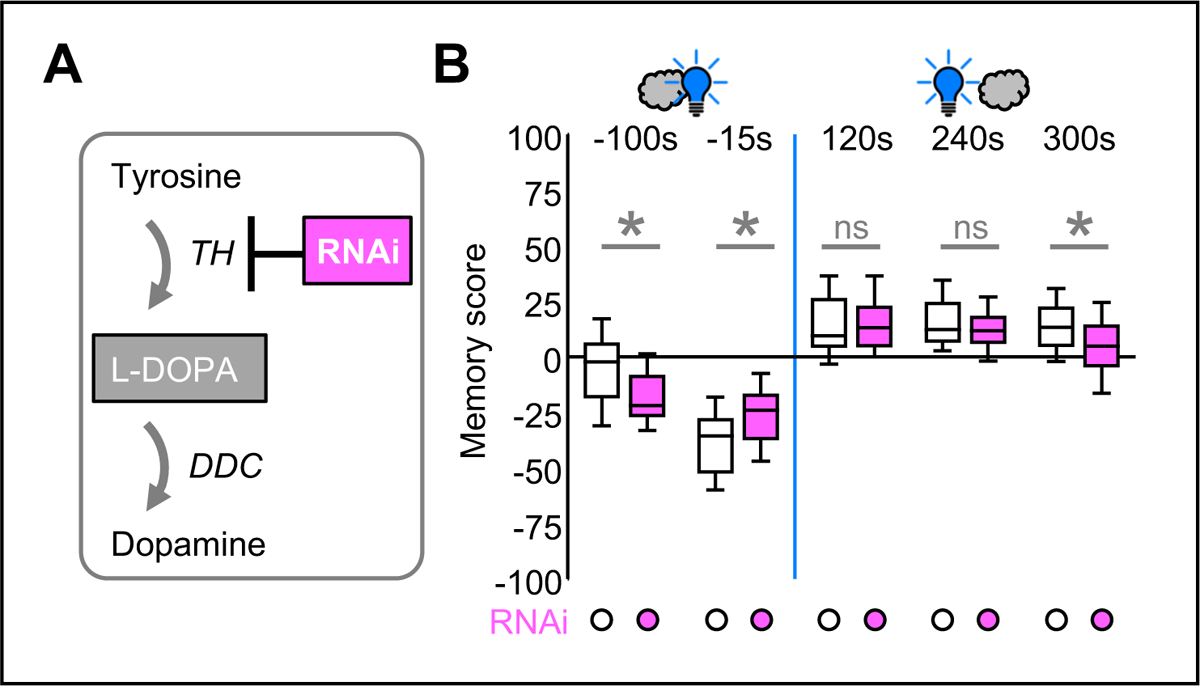
Local knock-down of the TH enzyme both extends and blunts the temporal profile of reinforcement by PPL1-01. **(A)** Schematic of dopamine biosynthesis and of the inhibition of tyrosine hydroxylase (TH) by RNA interference (RNAi). **(B)** Mapping out the effect of TH-RNAi in the PPL1-01 neuron on the temporal profile of PPL1-01 reinforcement (N= 32, 32; 34, 34; 40, 40; 33, 34; 42, 42). Relative to controls, TH-RNAi promotes punishment memory upon trace conditioning (ISI - 100s) and decreases punishment memory upon delay conditioning (ISI - 15s). Relief memory is decreased (ISI 300s). Control flies were of the genotype PPL1-01>ChR2-XXL (open box plots and circles); flies for TH knock-down in the PPL1-01 neuron additionally carried the TH-RNAi construct (PPL1-01>ChR2-XXL/TH-RNAi) (box plots and circles with magenta fill). Other details are as in the legend of Figure 1. * indicates significance in MW-U tests at an error rate of 5%, adjusted according to Bonferroni-Holm, “ns” indicates non-significance in such tests. Data and statistical results are documented in Supplementary Table 1; the underlying preference scores are shown in Supplementary Figure 6. Memory scores separated by sex are shown in Supplementary Figure 7.

We conclude that compromising TH function both extends and blunts the temporal profile of reinforcement by PPL1-01 (Figure 4A): trace conditioning (ISI - 100s) is improved, delay conditioning is impaired, and relief conditioning is abolished for longer intervals (ISIs 240s and 300s).

**Figure 4:**
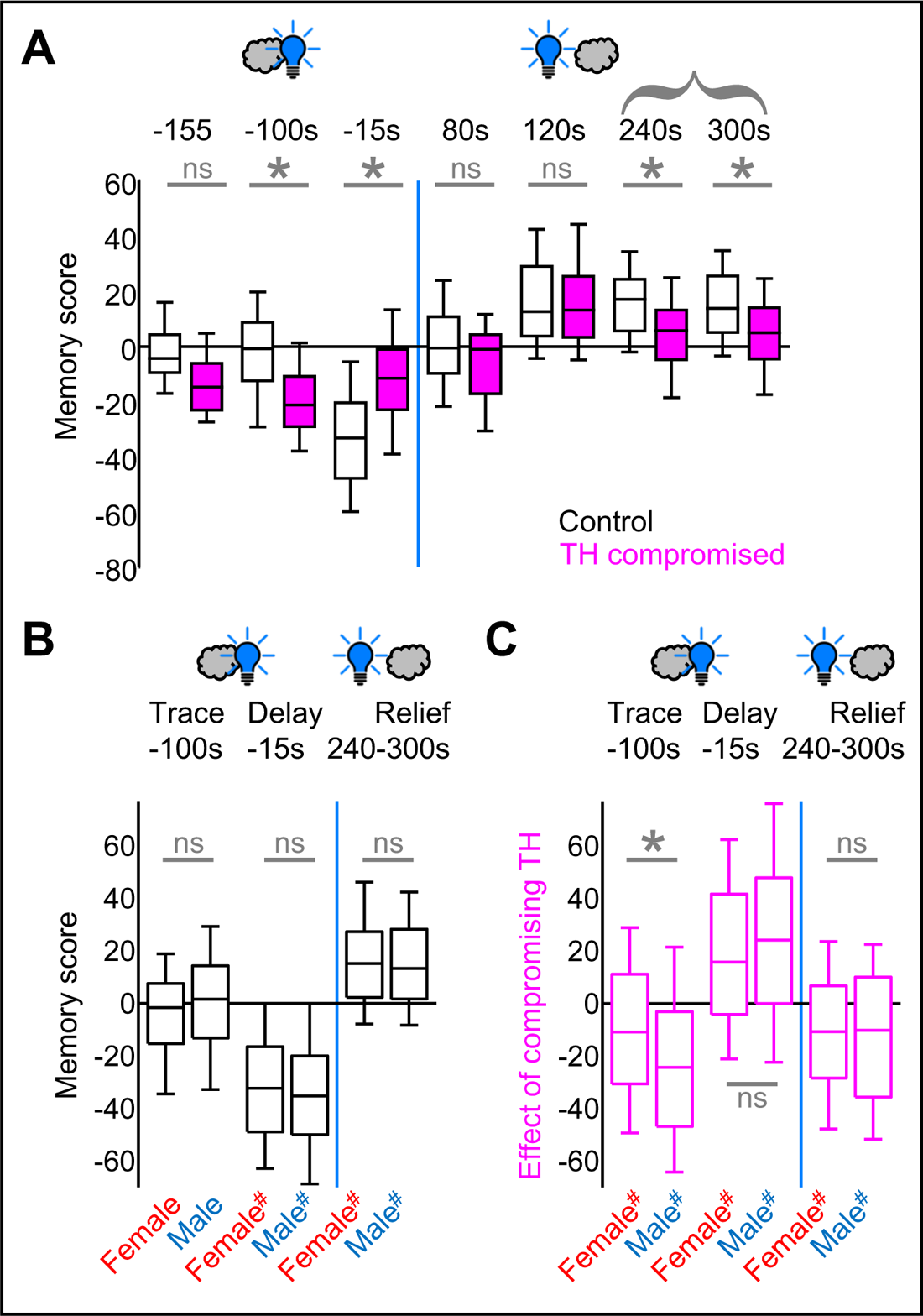
Compromising TH function both extends and blunts the temporal profile of reinforcement by PPL1-01. **(A)** Summary of the effects of compromising TH function on the temporal profile of PPL1-01 reinforcement, combined for 3IY and TH-RNAi, and across the present study. Shown are the memory scores of the respective control (open box plots) and TH-compromised cases (box plots and circles with magenta fill) (N= 20, 20, 120, 122, 159, 165, 20, 20, 101, 99, 108, 113, 62, 62). **(B)** Data for only the control cases shown in (A), separated by sex. Neither for trace conditioning (ISI - 100s), nor for delay conditioning (ISI - 15s), nor for relief conditioning (ISIs 240s and 300s) were sex-dependent differences observed (for a justification of why relief conditioning with ISIs of 80s and 120s is not included in this grouping see below) (N= 120, 120, 159, 157, 169, 169). **(C)** For the data in (A) the difference in memory scores of the TH-compromised cases minus the scores in the controls is plotted, separately for female and male flies, to quantify how strongly the compromising of TH function affects memory scores, in either sex. For trace conditioning (ISI - 100s), the effect of compromising TH function was less pronounced in females than in males, whereas no such difference was observed for delay (ISI - 15s) and relief conditioning (N= 118, 119, 159, 157, 168, 167). For relief conditioning, data were considered only for those ISIs for which the compromising of TH function was observed to have an effect to begin with (A: 240s and 300s). Other details are as in the legend of Figure 1. * indicates significance in MW-U tests at an error rate of 5%, adjusted according to Bonferroni-Holm, “ns” indicates non-significance in such tests. “#” indicates significance in OSS-tests at an error rate of 5%, adjusted according to Bonferroni-Holm. Data and statistical results are documented in Supplementary Table 1.

### Intelude: Separating data by sex

Our lab routinely acquires data separated by sex, but in aversive short-term memory we have so far never observed reliable differences between female and male flies. Accordingly, for the control conditions of the present study memory scores do not differ between the sexes for trace, delay, or relief conditioning (Figure 4B) (Supplementary Figure 3, Supplementary Figure 4, Supplementary Figure 7, Supplementary Figure 9). In ascertaining the difference in memory scores of the TH-compromised case minus the controls, however, we were surprised to find that for trace conditioning the effect of compromising the TH function, which was significant in both sexes, was selectively less pronounced in females than in males (Figure 4C). Two observations suggest that this sex difference is not a statistical artifact. Firstly, this sex difference can be discerned for both 3IY feeding as an acute, systemic intervention (Supplementary Figure 3C, Supplementary Figure 3D) and for TH-RNAi as a constitutive, cell-specific intervention (Supplementary Figure 7). Secondly, when the brain-wide HPLC measurements of biogenic amines after 3IY feeding were separated by sex, this revealed only a non-significant tendency toward a decrease in dopamine levels in the females, whereas a significant decrease in dopamine levels was observed only in the males (Supplementary Figure 5). This suggests that decreases in dopamine levels that in females remain below the significance threshold in brain-wide HPLC measurements (Supplementary Figure 5) can nevertheless have behavioral effects (Figure 4C), and that these behavioral effects are weaker in females than those produced by the more pronounced decreases in dopamine levels in males (Figure 4C, Supplementary Figure 5).

### An inhibitor of the TPH enzyme can reverse the effects of 3IY on trace conditioning

PPL1-01 is one of a total of 12-15 dopaminergic neurons in the PPL1 cluster, and specifically in the subset of six of these that innervate the mushroom body (Mao and Davis 2009; Aso et al. 2014; Li et al. 2020). It has been reported that within the PPL1 cluster there is at least one neuron that is not only immunoreactive against TH but also against serotonin, and that constitutively compromising TH function by genetic means can both increase the number of anti-serotonin immunoreactive neurons in the PPL1 cluster and alter patterns of anti-serotonin immunoreactivity in the mushroom body, specifically at the tips of the α and α’ lobes, which receive both dopaminergic and serotonergic input (Niens et al. 2017). Preliminary results suggested that our method for acutely lowering dopamine levels by 3IY left the number of anti-serotonin immunoreactive neurons in the PPL1 cluster unchanged but altered patterns of serotonin-immunoreactivity as previously reported (not shown). This encouraged us to test whether downregulating serotonin synthesis would alter the effects we observed by feeding 3IY. To test for this possibility, we used para-chlorophenylalanine (PCPA), an inhibitor of the tryptophan hydroxylase enzyme (TPH) (Figure 5A). For trace conditioning (ISI - 100s), the increase in punishment memory caused by 3IY was fully reversed by an additional feeding of PCPA (Figure 5B). For delay conditioning (ISI - 15s), the 3IY-induced decrease in punishment memory was only partially reversed by PCPA. For relief conditioning (ISI 240s), however, the decrease in relief memory through 3IY was not moderated by PCPA. Under low-dopamine conditions, the additional feeding of PCPA thus had a graded effect on memory scores, in the sense that it was strong for trace conditioning (ISI - 100s) and tapered off as the ISIs were increased to −15s and 240s. Of note is that feeding PCPA alone, that is feeding PCPA under conditions of normal dopamine levels, had no effect on either form of conditioning (Figure 5C).

**Figure 5:**
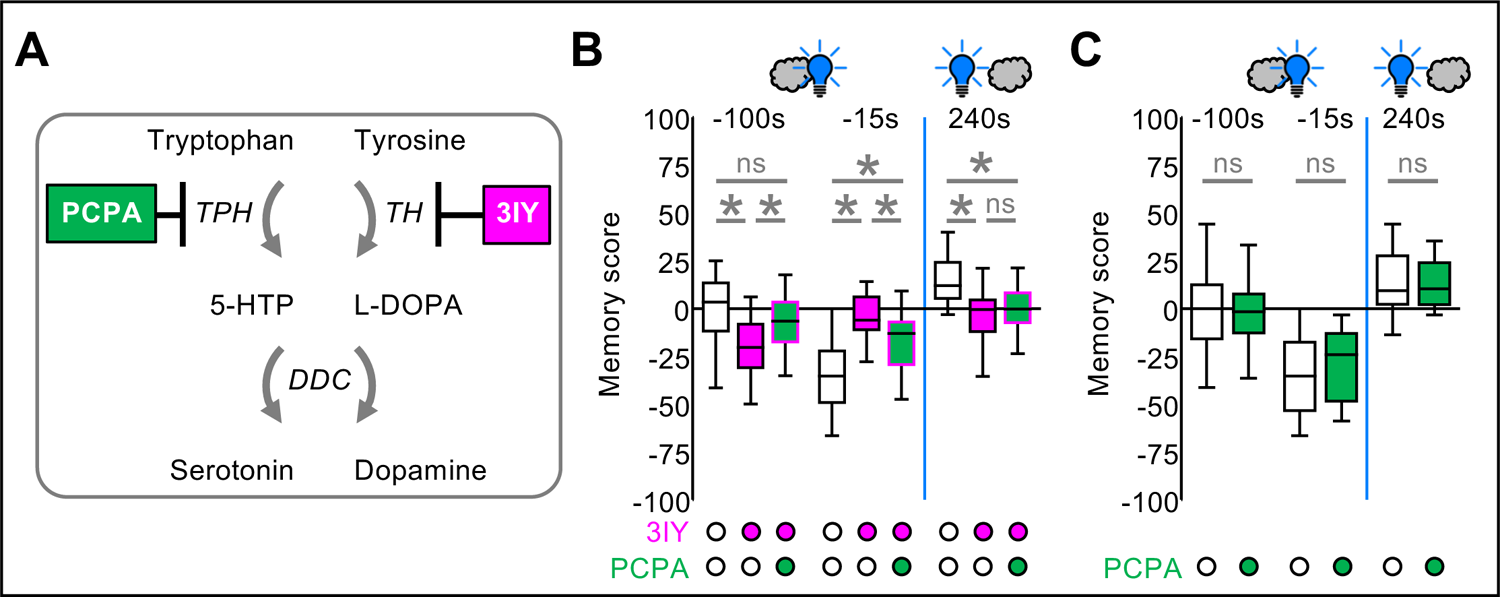
Temporal profile of reinforcement by PPL1-01 upon pharmacologically inhibiting the TPH and the TH enzyme. **(A)** Schematic of serotonin and dopamine biosynthesis, of the inhibition of tryptophan hydroxylase (TPH) by para-chlorophenylalanine (PCPA), and of the inhibition of tyrosine hydroxylase (TH) by 3-iodo-L-tyrosine (3IY), respectively. 5-HTTP: 5-hydroxytryptophan; L-DOPA: 3,4-dihydroxy-L-phenylalanine; DDC: dopamine decarboxylase. Drug feeding was performed by the food method. **(B)** Relative to controls, punishment memory after odor→PPL1-01 trace conditioning (ISI - 100s) is increased upon feeding of 3IY, an effect that is partially reversed by an additional feeding of PCPA (N= 38, 39, 45). For delay conditioning (ISI - 15s), punishment memory is reduced by 3IY, an effect that is partially reversed by PCPA (N= 29, 33, 32). The reduction of relief memory (ISI 240s) by 3IY was not reversed by PCPA (N=25, 28, 29). **(C)** PCPA alone has no effect on punishment memory after trace conditioning (ISI - 100s) (N= 39, 39) or after delay conditioning (ISI - 15s) (N= 45, 48) and leaves relief memory intact, too (ISI 240s) (N= 44, 41). Open box plots and circles refer to the control condition, magenta and green fill to groups fed with 3IY or with 3IY plus PCPA, respectively. Flies were of the genotype PPL1-01>ChR-2XXL. Other details are as in the legend of Figure 1. * indicates significance and “ns” non-significance in MW-U tests at an error rate of 5%, adjusted according to Bonferroni-Holm. Data and statistical results are documented in Supplementary Table 1; the underlying preference scores are shown in Supplementary Figure 8. Memory scores separated by sex are shown in Supplementary Figure 9.

## Discussion

We report that compromising TH function, either acutely by pharmacological means (Figure 1, Figure 5, Supplementary Figure 4) or by cell-specific RNAi (Figure 3), both extends and blunts the temporal profile of PPL1-01 reinforcement. Specifically, it improves trace conditioning (ISI - 100s), impairs delay conditioning (ISI - 15s), and abolishes relief conditioning for longer intervals (ISIs 240s and 300s) (Figure 4A).

### Relief conditioning with short versus long ISIs

The present results on relief conditioning can reconcile earlier reports by König et al. (2018, loc. cit. figure 5) and Aso et al. (2019, loc. cit. figure 5, supplementary figure 3C). These studies differ in a number of ways, making it difficult to quantitatively relate the ISIs that were used. Differences include the use of a two-arm T-maze versus a horizontal, 4-field arena setup, and the use of ChR2XXL versus CsChrimson-Venus as the optogenetic effector, respectively. Nonetheless, by mapping out multiple ISIs, the present results confirm the TH-independence reported by König et al. (2018) for short relief ISIs, whereas for longer relief ISIs they are consistent with the TH-dependence reported by Aso et al. (2019).

### Trace versus delay conditioning

Our results show a dissociation between trace and delay conditioning. Trace conditioning, surprisingly, is improved by compromising TH function, whereas as expected (König et al. 2018; Aso et al. 2019) delay conditioning is impaired (Figure 4A, ISIs - 100s versus −15s). This contrast adds to earlier evidence that trace and delay procedures engage partially distinct downstream mechanisms in odor-shock associative learning. Mutants in the rutabaga gene, coding for the type I adenylate cyclase which acts as a molecular coincidence detector for this association, are unaffected in trace conditioning but are impaired in delay conditioning (Shuai et al. 2011). In turn, expression of a dominant-negative form of the Rac protein improves trace conditioning but leaves delay conditioning unaffected (Shuai et al. 2011). In a visual learning paradigm, Grover et al. (2022) showed a selective role for the dopamine receptor Dop2R (CG33517) for trace but not delay conditioning.

Trace and delay conditioning certainly also have features in common. For odor-shock associations these commonalities include impairment in mutants lacking Synapsin (Niewalda et al. 2015), asymptotic memory strength upon repeated training trials (albeit reached at a slower rate for trace conditioning), their rate of memory decay, the profile of generalization (Galili et al. 2011), the likely site of the memory trace in the mushroom body, and their requirement of the dopamine receptor Dop1R1 (CG9652) (Shuai et al. 2011) (also see Grover et al. 2022).

### A role for serotonin in trace conditioning – under low-dopamine conditions

We show that pharmacologically compromising TH function addresses dopamine signaling, leading to a decrease in brain-wide levels of dopamine rather than serotonin (Figure 2). Further, both this decrease in dopamine levels and the abovementioned effects on reinforcement learning were rescued by L-DOPA (Figure 1G, Figure 2). Interestingly, under low-dopamine conditions our results uncover a role for serotonin that is particularly strong for trace conditioning (ISI - 100s), and that tapers off with increasing ISIs (Figure 5B). The fact that this role for serotonin is observed under conditions of normal and indeed tendentially lowered brain-wide serotonin levels (Figure 2) suggests that only relatively few neurons are mediating this effect. Under conditions of normal dopamine levels, however, our data do not provide evidence for a role of serotonin (Figure 5E); it is unclear whether this is at variance with the experiments reported by Zeng et al. (2023), as these lacked critical genetic and procedural controls.

### Implications of compromised TH function

At a cognitive level, one may view long-gap trace-conditioned cues as providing no evidence, and delay- and relief-conditioned cues as providing respectively evidence for and evidence against the causation of punishment. The temporal profile of reinforcement upon the compromising of TH function (Figure 4A) would thus imply a delusion-like state, in which causality is attributed to cues that do not merit it, whereas credible evidence in favor of as well as credible evidence against the causation of punishment is not properly appreciated. In a human subject, this would result in delusional beliefs about causal event structure, providing a hallmark symptom of schizophrenia (McCutcheon et al. 2019). Indeed, human schizophrenia patients tend to perceive cause-effect relationships where they do not exist and ‘jump to conclusions’ without sufficient evidence (Garety et al. 2005; Moritz and Woodward 2005; Uhlhaas and Silverstein 2005; Dudley et al. 2016). Behavior based on causal belief structures that are thus insufficiently grounded in experience would also lead to poor performance in various memory tasks. The implication is that a dysfunction in dopaminergic reinforcement processing, or possibly a distorted dopamine-serotonin balance, is a common endophenotype of both positive symptoms (delusional causal beliefs) and cognitive symptoms (impaired memory performance) in schizophrenia.

In practical terms, the present study shows that mapping out the full ‘fingerprint’ of reinforcement across multiple temporal intervals can be required to understand how a given manipulation affects reinforcement learning. Indeed, our results (Figure 4A) provide a case of an experimental manipulation where picking any one single interval between odor and reinforcement will lead to drastically different conclusions, namely that memory is improved, impaired, or unaffected, depending on which interval is chosen. It therefore seems possible that apparent discrepancies in the literature can be resolved through mapping the full temporal profile of reinforcement.

## Supporting information

Amin et al PPL1 bioRxiv Supplemental Figures

Amin et al PPL1 bioRxiv Supplementary Table 1

## Acknowledgements

Discussions with P. Stevenson (U Leipzig), M. Fendt (OVGU Magdeburg) and Juliane Thöner (LIN) are gratefully acknowledged. We thank R.D.V. Glasgow (Zaragoza, Spain) for language editing.

