## Supplementary material for "Compromising tyrosine hydroxylase function establishes a delusion-like temporal profile of reinforcement by dopamine neurons in *Drosophila*": Amin et al PPL1 bioRxiv Supplemental Figures

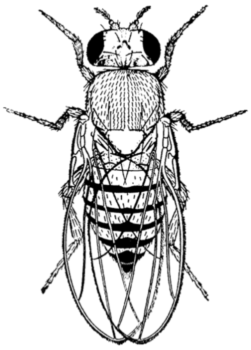

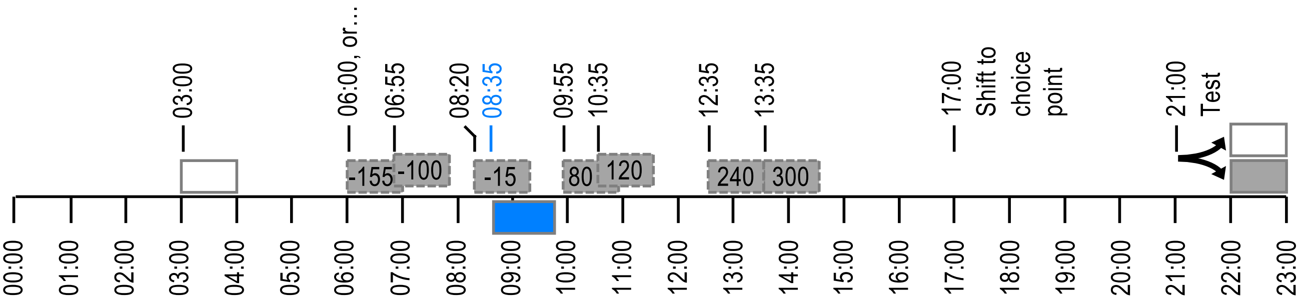


**Supplementary Figure 1: Timing of pairing odor and PPL1-01 activation by blue light**

At time 0:00 min, the flies were gently loaded into the experimental setup. From 3:00 min on, the reference odor (white box) was presented for 1 min. For the optogenetic activation of the PPL1-01 neuron, blue light (blue box) was applied at 8:35 min for 1min with the help of 24 LEDs of 465 nm peak wavelength mounted on the inner surface of 2.5 cm-diameter and 4.5 cm-length hollow cylinders. These cylinders were fitted around transparent training tubes harboring the flies. Blue light was applied as 12 pulses, each 1.2-sec long and followed by the next pulse with a 5 sec onset-to-onset interval. The absolute irradiance in the middle of the training tube during blue light pulses was 200 µW/cm$^2$ as measured with an STS-VIS Spectrometer (Ocean Optics). The paired odor (grey box) was presented for 1 min, too, at the onset-to-onset inter-stimulus-intervals (ISIs) from the blue light as indicated by the numbers within the grey boxes (s). Negative ISI values indicate that the presentation of the paired odor started before the blue light (ISIs -155s, -100s and -15s); positive ISIs indicate the reverse order of events (ISIs 80s, 120s, 240s and 300s). Onset times for all ISIs are indicated above. At 17:00 min the flies were shifted to the choice point between the odors. At 21:00 min the test started and the flies were released into the T-maze. After 2 min the arms of the maze were closed and the flies on each side were counted.

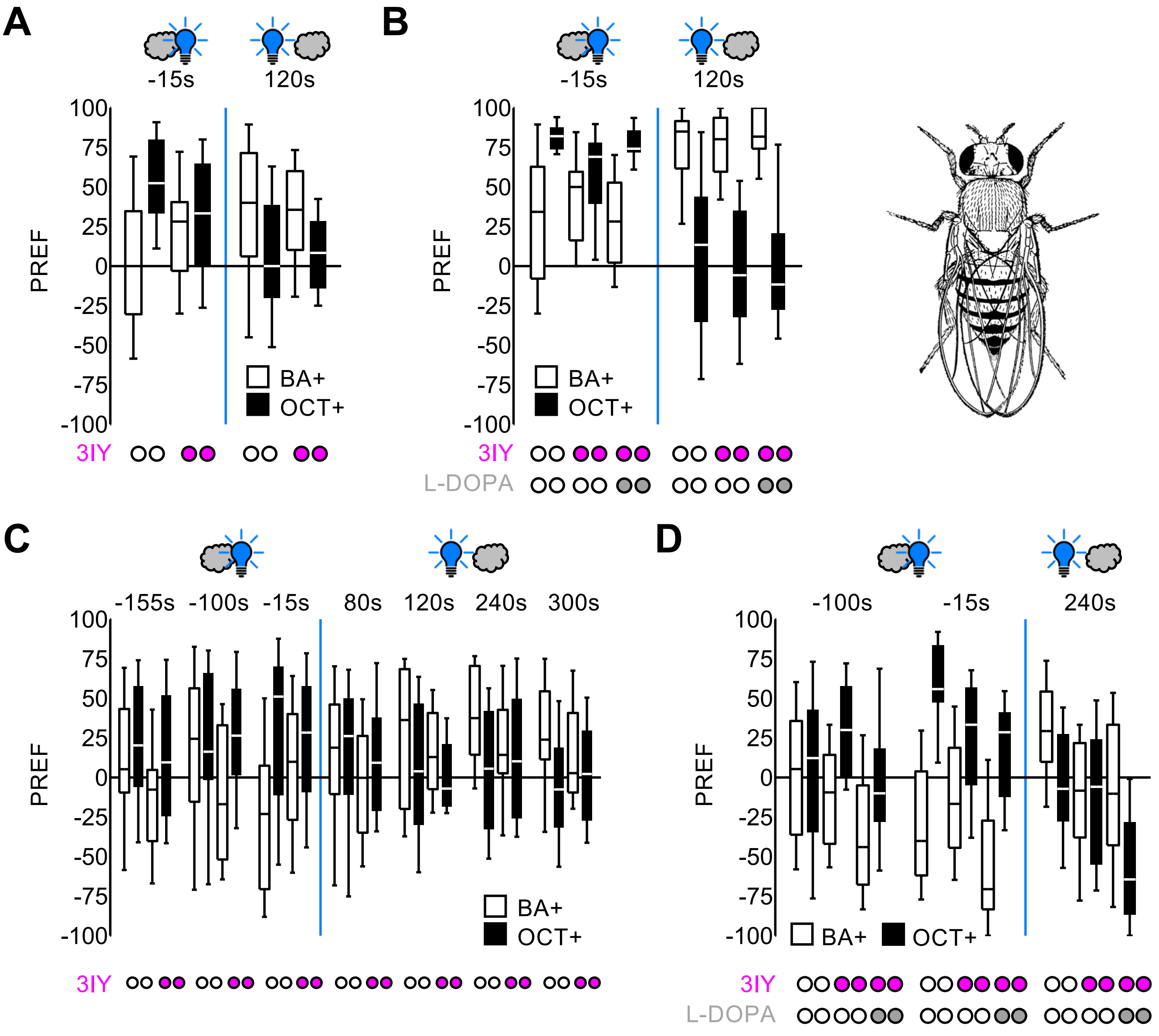


**Supplementary Figure 2: Preference scores underlying the memory scores in Figure 1**

**(A-D)** Preference scores calculated from the choice between the odors benzaldehyde and octanol (BA, OCT) according to equation 1, and as underlying the memory scores in Figure 1D-G, respectively. Box plots represent the median as the middle line, 25%/75% quantiles as box boundaries, and 10%/90% quantiles as whiskers. Open box plots show benzaldehyde preference after BA was paired with PPL1-01 activation and OCT served as reference odor (BA+), black fill indicates BA preference after OCT was paired with PPL1-01 activation and BA served as a reference odor (OCT+). The grey clouds depict the paired odor, and the blue light bulb PPL1-01 activation, presented with the indicated temporal relationship. Open circles below the panels refer to the control conditions, magenta fill to the 3IY-fed cases, grey fill to those additionally fed with L-DOPA. Other details as in the legend of Figure 1. Data are documented in Supplementary Table 1.

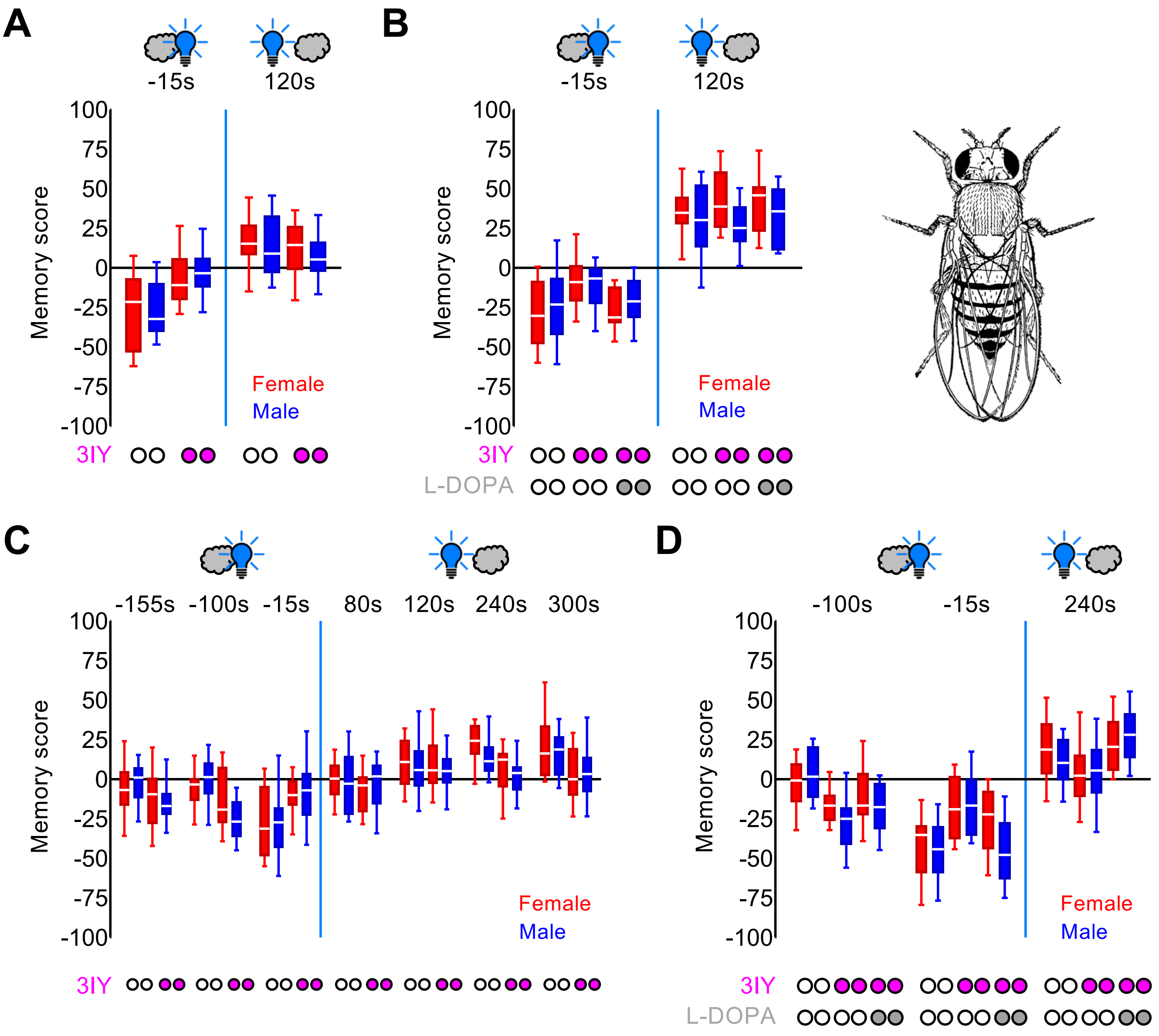


**Supplementary Figure 3: Memory scores from Figure 1 separated by sex**
**(A-D)** Memory scores from Figure 1D-G, respectively, separated by sex. Box plots represent the median as the middle line, 25%/75% quantiles as box boundaries, and 10%/90% quantiles as whiskers. Red fill of the box plots shows data from females, blue fill indicates data from males. The grey clouds depict the paired odor, and the blue light bulb PPL1-01 activation, presented with the indicated temporal relationship. Open circles below the panels refer to the control conditions, magenta fill to the 3IY-fed cases, grey fill to those additionally fed with L-DOPA. Other details as in the legend of Figure 1. Data are documented in Supplementary Table 1.

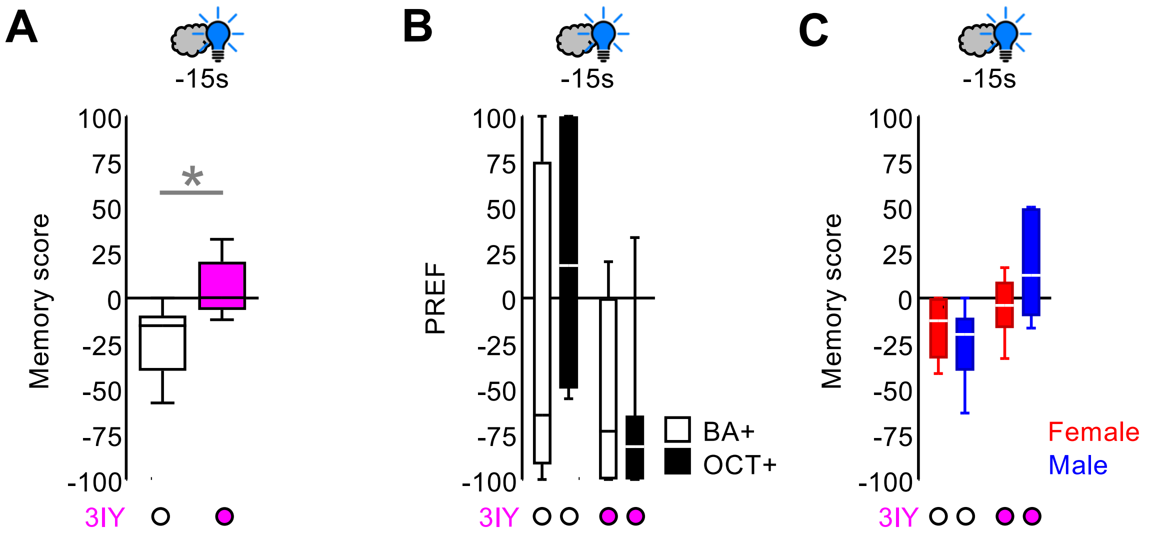


**Supplementary Figure 4: Repetition of the experiment in Figure 1D (ISI -15s)**

**(A-C)** Repetition of the experiment shown in Figure 1D, for the ISI of -15s. Box plots represent the median as the middle line, 25%/75% quantiles as box boundaries, and 10%/90% quantiles as whiskers. The grey clouds depict the paired odor, and the blue light bulb PPL1-01 activation, presented with the indicated temporal relationship. (A) Memory scores, determined according to equation 2, for the control condition (open box plot, N= 6) and the 3IY-fed case (magenta fill, N= 6). * indicates significance in a MW-U test. (B) Preference scores, calculated according to equation 1, underlying the data from (A). Open box plots show benzaldehyde preference after BA was paired with PPL1-01 activation and OCT served as a reference odor (BA+); black fill indicates BA preference after OCT was paired with PPL1-01 activation and BA served as a reference odor (OCT+). (C) Data from (A), separated by sex. Red fill of the box plots refers to data from females, blue fill to data from males. Open circles below the panels refer to the control conditions, magenta fill to the 3IY-fed cases. Other details as in the legend of Figure 1. Data are documented in Supplementary Table 1.

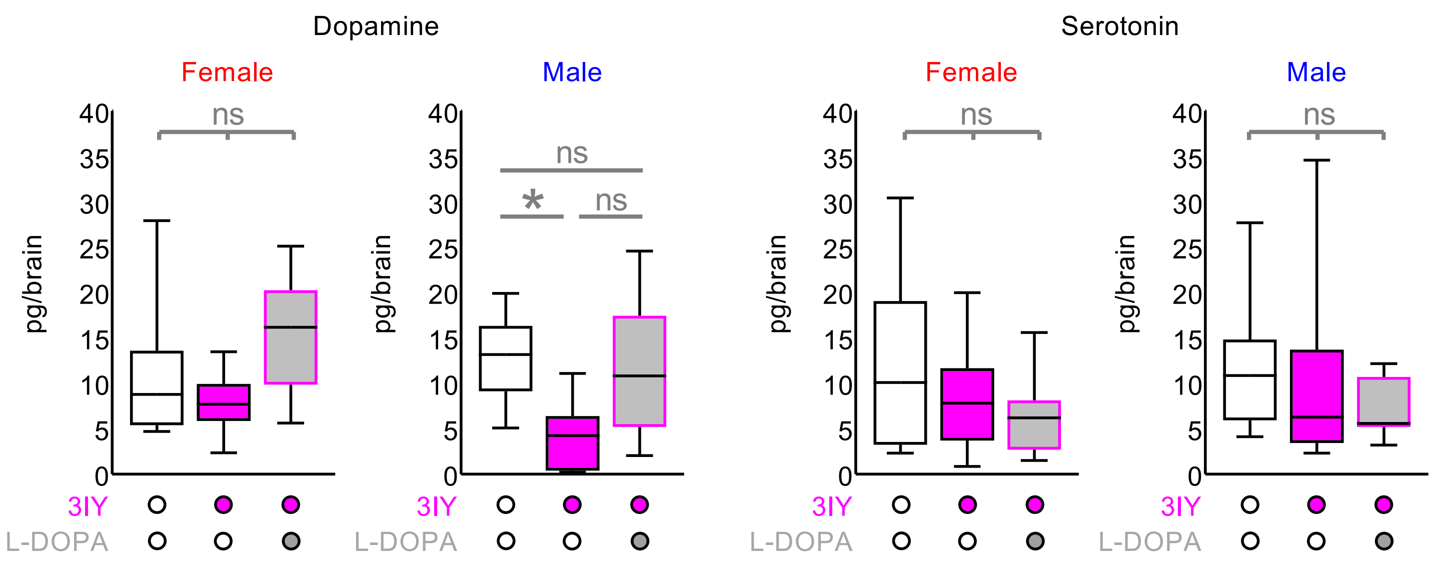


**Supplementary Figure 5: Levels of biogenic amines from Figure 2 separated by sex**
Brain-wide levels of dopamine (left) and serotonin (right) from Figure 2, separated for females and males as indicated. Box plots represent the median as the middle line, 25%/75% quantiles as box boundaries, and 10%/90% quantiles as whiskers. Open box plots and circles refer to the control condition, magenta and grey fill to groups fed with 3IY or with 3IY plus L-DOPA, respectively.
"ns" indicates non-significance in KW-tests, except for the case of dopamine measurements in males where * and "ns" refer to significance and non-significance, respectively, in MW-U tests at an error rate of 5%, adjusted according to Bonferroni-Holm. Other details as in the legend of Figure 2. Data are documented in Supplementary Table 1.

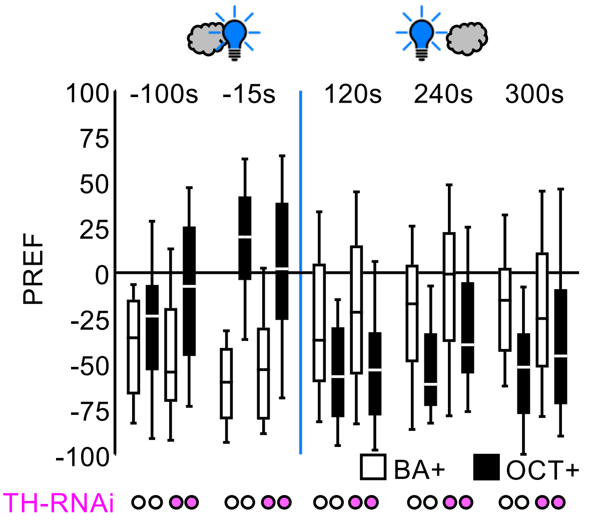


**Supplementary Figure 6: Preference scores underlying the memory scores in Figure 3**
Preference scores calculated from the choice between the odors benzaldehyde and octanol (BA, OCT) according to equation 1, and as underlying the memory scores in Figure 3B. Box plots represent the median as the middle line, 25%/75% quantiles as box boundaries, and 10%/90% quantiles as whiskers. Open box plots show benzaldehyde preference after BA was paired with PPL1-01 activation and OCT served as a reference odor (BA+); black fill indicates BA preference after OCT was paired with PPL1-01 activation and BA served as a reference odor (OCT+). The grey clouds depict the paired odor and the blue light bulb PPL1-01 activation, presented with the indicated temporal relationship. Open circles below the panels refer to the control conditions, magenta fill to the cases with RNAi in the TH neurons. Other details as in the legend of Figure 1. Data are documented in Supplementary Table 1.

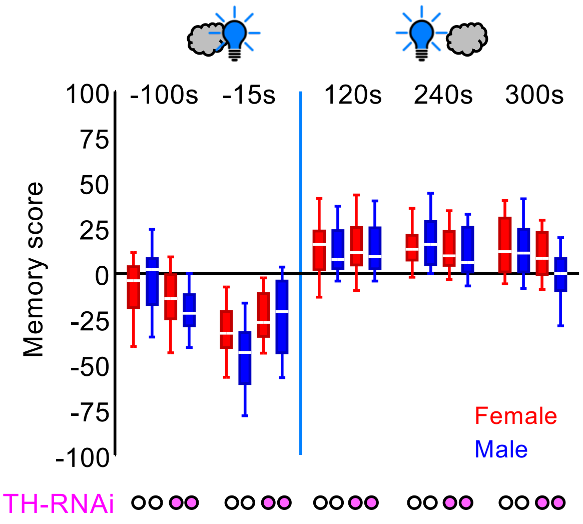


**Supplementary Figure 7: Memory scores from Figure 3 separated by sex**

Memory scores from Figure 3B, separated by sex. Box plots represent the median as the middle line, 25%/75% quantiles as box boundaries, and 10%/90% quantiles as whiskers. Box plots with red fill show data from females, blue fill indicates data from males. The grey cloud depicts the paired odor, and the blue light bulb PPL1-01 activation, presented with the indicated temporal relationship. Open circles below the panels refer to the control conditions, magenta fill to the cases with RNAi in the TH neurons. Other details as in the legend of Figure 1. Data are documented in Supplementary Table 1.

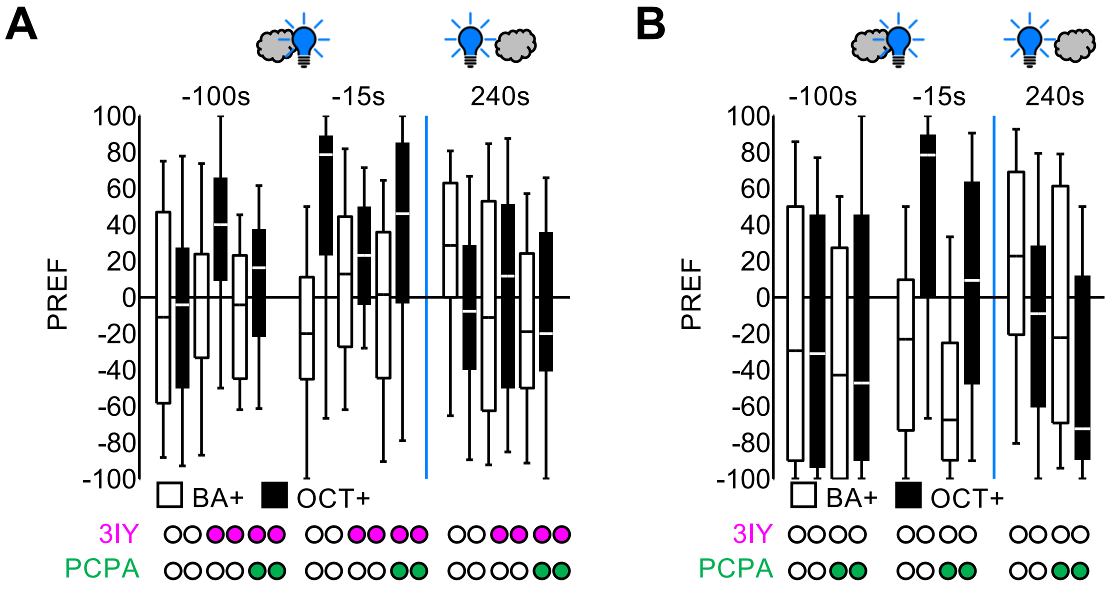


**Supplementary Figure 8: Preference scores underlying the memory scores in Figure 5**
**(A and B)** Preference scores calculated from the choice between the odors benzaldehyde and octanol (BA, OCT) according to equation 1, and as underlying the memory scores in Figure 5B and C, respectively. Box plots represent the median as the middle line, 25%/75% quantiles as box boundaries, and 10%/90% quantiles as whiskers. Open box plots show benzaldehyde preference after BA was paired with PPL1-01 activation and OCT served as a reference odor (BA+), black fill indicates BA preference after OCT was paired with PPL1-01 activation and BA served as a reference odor (OCT+). The grey clouds depict the paired odor, and the blue light bulb PPL1-01 activation, presented with the indicated temporal relationship. Open circles below the panels refer to the control conditions, magenta fill refers to feeding with 3IY, and green fill to feeding with PCPA in addition. Other details as in the legend of Figure 1. Data are documented in Supplementary Table 1.

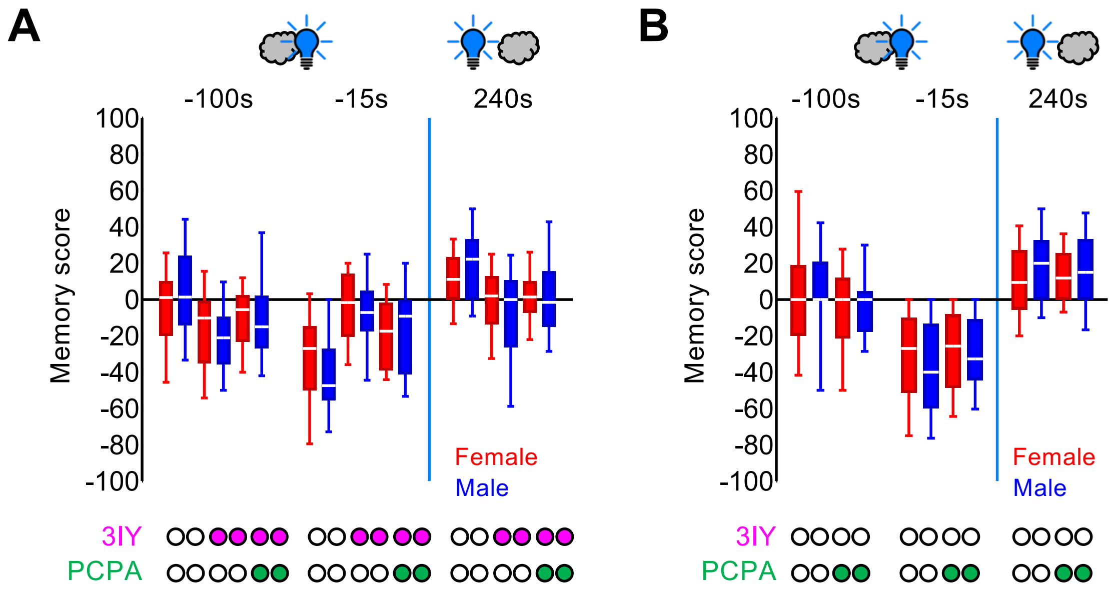


**Supplementary Figure 9: Memory scores from Figure 5 separated by sex**
**(A and B)** Memory scores from Figure 5B and C, respectively, separated by sex. Box plots represent the median as the middle line, 25%/75% quantiles as box boundaries, and 10%/90% quantiles as whiskers. Red fill of the box plots shows data from females, blue fill indicates data from males. The grey clouds depict the paired odor, and the blue light bulb PPL1-01 activation, presented with the indicated temporal relationship. Open circles below the panels refer to the control conditions, magenta fill refers to feeding with 3IY, and green fill to feeding with PCPA in addition. Other details as in the legend of Figure 1. Data are documented in Supplementary Table 1.
